# ROCK1 activates mitochondrial fission leading to oxidative stress and muscle atrophy

**DOI:** 10.1101/2023.10.22.563469

**Authors:** Meijun Si, Rizhen Yu, Hongchun Lin, Feng Li, Sungyun Jung, Sandhya S. Thomas, Farhard S Danesh, Yanlin Wang, Hui Peng, Zhaoyong Hu

**Author notes:** Meijun Si, Rizhen Yu and Hongchun Lin contributed equally to this work. **Correspondence:** Dr. Zhaoyong Hu: Nephrology Division, Department of Medicine, Baylor College of Medicine, ABBR704, One Baylor plaza, Houston, TX 77030. Hui Peng, Nephrology Division, Third Affiliated Hospital of Sun Yat-sen University, 600 Tianhe road, Guangzhou, China.

## Abstract

Chronic kidney disease (CKD) is often associated with protein-energy wasting (PEW), which is characterized by a reduction in muscle mass and strength. Although mitochondrial dysfunction and oxidative stress have been implicated to play a role in the pathogenesis of muscle wasting, the underlying mechanisms remain unclear. In this study, we used transcriptomics, metabolomics analyses and mouse gene manipulating approaches to investigate the effects of mitochondrial plasticity and oxidative stress on muscle wasting in mouse CKD models. Our results showed that the expression of oxidative stress response genes was increased, and that of oxidative phosphorylation genes was decreased in the muscles of mice with CKD. This was accompanied by reduced oxygen consumption rates, decreased levels of mitochondrial electron transport chain proteins, and increased cellular oxidative damage. Excessive mitochondrial fission was also observed, and we found that the activation of ROCK1 was responsible for this process. Inducible expression of muscle-specific constitutively active ROCK1 (mROCK1^ca^) exacerbated mitochondrial fragmentation and muscle wasting in CKD mice. Conversely, ROCK1 depletion (ROCK1-/-) alleviated these phenomena. Mechanistically, ROCK1 activation promoted the recruitment of Drp1 to mitochondria, thereby facilitating fragmentation. Notably, the pharmacological inhibition of ROCK1 mitigated muscle wasting by suppressing mitochondrial fission and oxidative stress. Our findings demonstrate that ROCK1 participates in CKD-induced muscle wasting by promoting mitochondrial fission and oxidative stress, and pharmacological suppression of ROCK1 could be a therapeutic strategy for combating muscle wasting in CKD conditions.

**Translational Statement:** Protein-energy wasting (PEW) is a prevalent issue among patients with chronic kidney disease (CKD) and is characterized by the loss of muscle mass. Our research uncovers a critical role that ROCK1 activation plays in muscle wasting induced by CKD. We found that ROCK1 is instrumental in causing mitochondrial fission, which leads to increased oxidative stress in muscle cells. By employing a pharmacological inhibitor, hydroxyfasudil, we were able to effectively curb ROCK1 activity, which in turn mitigated muscle wasting by reducing both mitochondrial fission and oxidative stress. These findings suggest that pharmacological inhibition of ROCK1 presents a promising therapeutic strategy for combating the muscle wasting associated with CKD.

## Introduction

Protein-energy wasting is a prevalent syndrome associated with several catabolic diseases such as chronic kidney disease (CKD). Its clinical manifestation includes the loss of skeletal muscle mass, decreased muscle strength, and poor quality of life, which ultimately increases morbidity and mortality ^1, 2^. Although the molecular mechanisms involved in muscle wasting are not yet fully understood, it is believed to be a multifactorial process leading to the disruption of muscle homeostasis ^3^. It involves increased proteolysis via the ubiquitin-proteasome system (UPS) and the autophagosome/lysosome, decreased protein synthesis, or a combination of both ^4, 5^. Various biological events have been implicated in the development of muscle catabolism, including decreased anabolic factors like growth hormone ^6, 7^, increased catabolic cytokines such as tumor necrosis factor alpha (TNF-α) and TNF-related weak inducer of apoptosis (TWEAK) ^8, 9^, and increased reactive oxygen species (ROS) emission from malfunctioning mitochondria ^3, 10^. These events can occur simultaneously, leading to muscle catabolism, as observed in CKD, where inflammation and suppressed insulin-like growth factor (IGF)-1/insulin signaling promote muscle protein degradation through UPS and induce muscle wasting ^11^.

Skeletal muscles depend on their mitochondrial networks for energy production, where oxidative phosphorylation (OXPHOS) and β-oxidation take place to generate ATP. However, the mitochondrial network is also responsible for producing approximately 90% of cellular ROS during this process ^12^. Therefore, if the mitochondrial network is damaged, not only will muscle oxidative capacity and ATP production be reduced, but mitochondrial ROS emissions might also increase. When cellular ROS levels exceed the antioxidant capacity of a cell, these oxidants may modify key cellular molecules such as proteins, DNA, and lipids, which can eventually lead to cellular injury and the suppression of protein synthesis ^13^. Indeed, previous studies have suggested that excessive mitochondrial ROS emissions may initiate the development of muscle atrophy ^14, 15^.

Rho-associated protein kinase-1 (ROCK1) is a downstream effector of small G proteins, such as RhoA, and plays essential roles in cell contraction, migration, cell cycle progression, and cellular metabolism ^16^. Additionally, ROCK1 can be constitutively activated by caspase-3, which is believed to mediate stress fiber formation, ROS production, and apoptosis ^17, 18^. Our previous study demonstrated that ROCK1 was activated in muscles of CKD mice, leading to suppression of p-Akt signaling and accelerated protein degradation ^19^. As ROCK1 also reportedly regulates cell energy metabolism by promoting mitochondrial fission in stem and epithelial cells ^20, 21^, we hypothesize that ROCK1 activation may trigger excessive mitochondrial fragmentation contributing CKD-induced muscle atrophy.

To test this possibility, we performed RNA sequencing (RNA-seq) on isolated tibialis anterior (TA) muscles from mice with CKD and analyzed the relationship between OXPHOS and oxidative stress during muscle wasting. We also investigated whether ROCK1 plays a key role in atrophic muscle by triggering mitochondrial fission. Finally, we tested whether the pharmacological inhibition of ROCK1 attenuated muscle protein loss in CKD mice. Our results identified ROCK1 as an upstream signaling molecule that promotes mitochondrial fission and oxidative stress and demonstrated that ROCK1 could be a potential intervention target for limiting the development of CKD-induced muscle wasting.

## Results

### Differential regulation of OXPHOS capacity and oxidative stress in muscles of mice with CKD

To investigate the transcript-level changes associated with muscle wasting in CKD, we conducted RNA sequencing (RNA-seq) analysis of the tibialis anterior (TA) muscles of mice with CKD. We defined muscle wasting as a decrease in the TA muscle weight/tibia length ratio of more than 20% compared to normal controls (Figure 1a), the presence of myofiber atrophy (Figure 1b and Supplementary Figure S1A), an obvious leftward shift in cross-sectional area (Figure 1c) and the upregulation of two well-known atrophy-related genes, Atrogin-1/Fbxo32 and MuRF-1/Trim63 (Figure 1d). We identified genes through RNA-seq analysis that exhibited significant changes in expression in the TA muscles of CKD mice, showing at least a 50% change (fold change > 1.5) in their expression values and having corrected p-values of less than 0.05 (Figure 1e). Using gene ontology analysis, we further identified several biological processes and pathways involved in muscle wasting, including protein degradation, autophagy, inflammation, energy metabolism, and oxidative stress (Figure 1f and Supplementary Figure S1B). Notably, the expression of genes involved in OXPHOS pathways was downregulated (Figure 1g), whereas the expression of oxidative stress-responsive genes was upregulated in the muscles of CKD mice (Figure 1h). We listed the top ten downregulated genes involved in OXPHOS pathways and the top ten upregulated oxidative stress-responsive genes. Gene set enrichment analysis (GSEA) confirmed the downregulation of the OXPHOS gene set and upregulation of the oxidative stress-responsive gene set in the muscles of mice with CKD (Figure 1i and j). The results suggest that OXPHOS reactions were hindered while oxidative stress was elevated in the muscles of CKD mice. The disparate regulation of these two pathways appears counterintuitive, as the main instigators of oxidative stress, cellular reactive oxygen species (ROS), are predominantly produced by OXPHOS reactions in the mitochondria ^22^. To decipher this paradox, we delved deeper into the underlying mechanism causing the heightened oxidative stress in the muscles of mice with CKD. We hypothesize that compromised mitochondrial integrity could be the key to understanding the distinct regulation of these two pathways.

**Fig.1.**
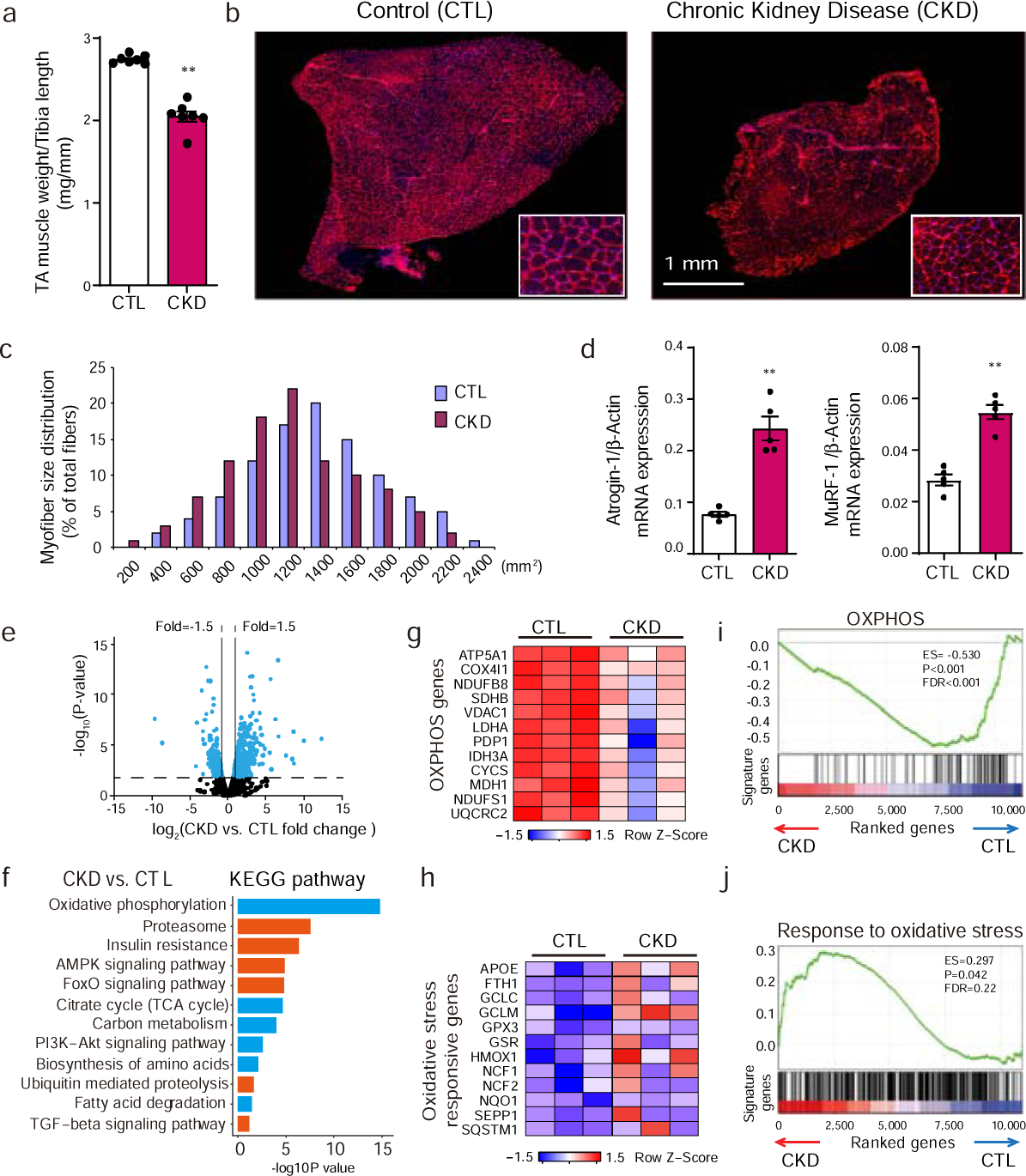
RNA-sequencing revealed suppressed OXPHOS and augmented oxidative stress in muscles of mice with CKD. (A) The quantification of tibialis anterior (TA) muscle weight normalized with the tibia length. Data were represented as mean±SEM, \*\**p*<0.01 vs. CTL (n = 9). (B) The cross-sectional area of cryosections (6μm) of TA muscle from mouse models of CKD and control mice are shown by immunostaining with anti-laminin antibody. Red color: laminin. Scale bars: 1mm. (C) The distribution of myofiber cross-sectional area (CSA) was shifted leftward in TA muscles from CKD mice compared with control mice (n = 5). (D) qRT-PCR confirms the upregulation of MuRF-1 and Atrogin-1 in the three muscle wasting models. Data are shown as means ± SEM, \*\**P* < 0.01, vs. Control (CTL), n = 5. (E) Volcano plot shows the results of RNA-sequencing (CKD vs. CTL, n=3 in each group). The dotted horizontal line represents *P*-value by *t*-test of 0.05. The dotted vertical lines indicate 1.5-fold change up and down respectively. (F) KEGG analysis of differentially expressed genes revealed by RNA-sequence of muscles from CKD mice compared with controls. Orange color indicates gene sets or pathways derived from genes that are significantly increased in catabolic models, while blue indicates results analyzed from genes that are significantly decreased in CKD muscles. (G and H) Heatmap shows expression of genes involved in OXPHOS pathways (G) and oxidative stress responsive genes (H) in the muscles from CKD or control mice. Color indicates the gene expression value scaled by Z score. (I and J) Gene Set Enrichment Analysis (GSEA) plots demonstrate enrichment score (ES) of gene sets in the RNA-sequencing data of muscles. Genes in each gene set are ranked by signal/noise ratio according to their differential expression between muscles form CKD and from control mice. The enrichment of oxidative phosphorylation (OXPHOS) gene set was significantly decreased in muscles of CKD mice (I), while gene set of response to oxidative stress was significantly enriched in CKD muscles (J).

### Excessive mitochondrial fission contributes to oxidative damage in muscles of mice with CKD

Mitochondria play a crucial role in consuming oxygen, primarily through the oxidative phosphorylation (OXPHOS) pathways. Impairment of these pathways can lead to a reduction in the oxygen consumption of myofibers. To investigate this further, we measured the cellular oxygen consumption rate (OCR) in freshly isolated rear flexor digitorum brevis (FBD) muscle fibers obtained from CKD and control mice. We also evaluated mitochondrial OXPHOS capacity by sequentially adding oligomycin, carbonyl cyanide-p-trifluoromethoxy phenylhydrazone (FCCP), and rotenone after measuring the basal OCR (Figure 2a). We found that basal OCR values were significantly suppressed in myofibers of CKD mice compared to control mice. Furthermore, proton leak values, which are linked to basic metabolic rate and body temperature, were increased in CKD myofibers. Additionally, maximal respiration rates, spare respiratory capacities, and ATP-related respiration were lower in myofibers from CKD mice than in those from control mice, indicating that oxidative metabolism was impaired in CKD myofibers (Figure 2a and Supplementary Figure S1C). Concurrently, the protein expression levels of respiratory electron transport chain subunits were decreased in muscles of CKD mice (Figure 2b), and succinate dehydrogenase (SDH) activities were also reduced (Figure 2c). These results confirmed the decline of oxidative metabolism in CKD muscles. In the context of low oxidative metabolism, CKD muscles were characterized by higher levels of protein carbonylation, indicating increased oxidative stress (Figure 2d). Metabolomics analysis showed that oxidative stress-related metabolites (such as tauropine, hypoxanthine, and cADP-ribose) were increased, while taurine, an antioxidant molecule ^14, 23, 24^, was reduced in muscles undergoing atrophy compared to normal muscles (Figure 2e, yellow bars). We also detected intermediate metabolites of glycolysis and fatty acid oxidation, which were significantly reduced in atrophic muscles compared to control muscles (Figure 2e, green and red bars). Using PhAM^excised^ Dentra-2 transgenic mice, in which mitochondria are labeled with a fluorescent protein ^25^, we found that excessive mitochondrial fission occurred during muscle wasting (Figure 2f). Electron microscopy confirmed the presence of smaller mitochondria or decreased mitochondrial density, along with a greater abundance of swollen mitochondria with disrupted cristae (Figure 2g), indicating that excessive mitochondrial fission was associated with mitochondrial damage. Taken together, these findings confirm the occurrence of significant mitochondrial impairments during CKD-induced skeletal muscle atrophy, which are associated with excessive mitochondrial fission and oxidative stress. Our next aim is to investigate the potential signaling pathways that promote mitochondrial fission during CKD-induced muscle wasting.

**Fig. 2.**
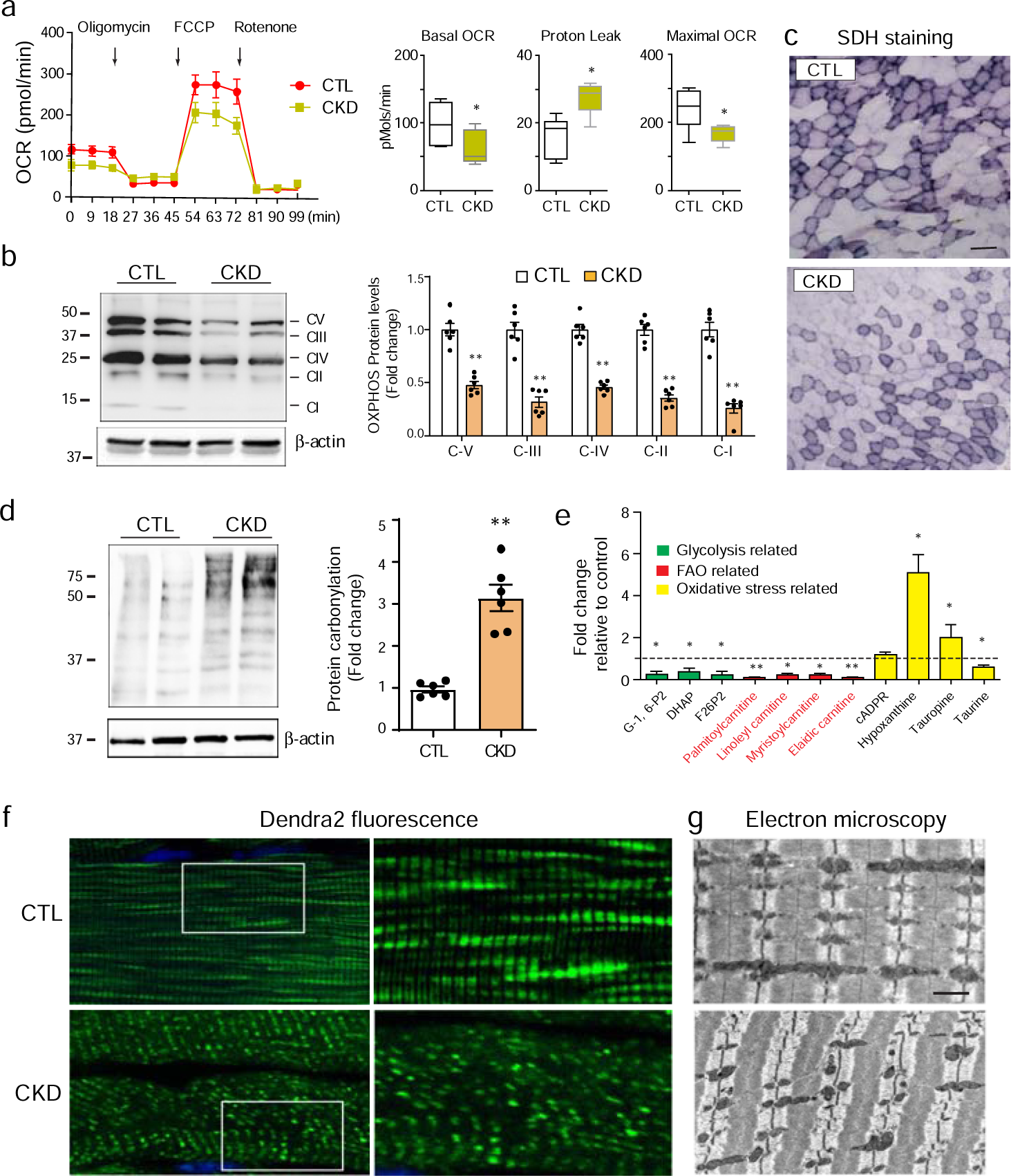
Impaired oxidative metabolism, augmented oxidative stress and mitochondrial fission are present simultaneously in atrophic muscles from CKD mice. (A) Cellular respiration of freshly isolated rear FDB myofibers was assessed by mito-stress test. The basal OCR, proton leak and maximal OCR were then calculated. All results were normalized to protein content and presented as mean ± SEM. (\**P*<0.05 vs. CTL, n=6). (B) Representative western blot shows the expression of subunits of mitochondrial respiratory electron transport chain in atrophying muscles (data are presented as mean ±SEM, \**P*<0.05, \*\**P*<0.01 vs. CTL, n=6). (C) Succinate dehydrogenase (SDH) activities were analyzed in sections of TA muscles from CKD mice. (D) Representative western blot demonstrated carbonyl levels of muscles from CKD mice by DNP antibody (data are presented as mean ±SEM, \**P*<0.05, \*\**P*<0.01 vs. CTL, n=6). (E) Metabolomics analysis of metabolites in TA muscles from CKD mice compared to the result of control mice. Selected metabolites were grouped in glycolysis (green), fatty acid oxidation (FAO, red) and oxidative stress (yellow). Data are presented as mean ± SEM. (\**P*<0.05, \*\**P*<0.01, n=5). (F) Representative microscopy images show fragmented mitochondria in CKD muscles of transgenic mice carrying Dendra-2 driven by Cox 8 promoter comparing with control (CTL) Dentra-2 transgenic mice. Zoomed areas are presented on the right panel (scale bar=10µm). (G) Transmission electron microscopy demonstrates the changes of mitochondria in sagittal sections of muscles from mice with CKD comparing with control mice (Scale bar = 500 nm).

### ROCK1 stimulates mitochondria fission resulting in oxidant stress in muscle cells

As a protein kinase, ROCK1’s activity is dictated by post-translational modifications rather than its transcription levels. There are two primary mechanisms that can activate ROCK1: through binding to RhoA (canonical activation) or via caspase-3 (constitutive activation), which cleaves the caspase-activated domain (CD domain) of ROCK1. In our previous study, we discovered that CKD activates ROCK1 in skeletal muscles through a caspase-3-mediated removal of the CD from ROCK1, without affecting the transcription levels of ROCK1 ^19^. Given that active ROCK1 induces mitochondrial fission in podocytes and endothelial cells ^21^, we hypothesized that the activation of ROCK1 leads to excessive mitochondrial fission in CKD muscles. To test this hypothesis, we initially examined whether overexpression of ROCK1 could trigger mitochondrial fragmentation in C2C12 myotubes. After transfecting C2C12 myotubes with an adenovirus carrying mouse full-length Rock1 for 48 hours, we evaluated the protein levels of both intact and constitutively activated ROCK1 (approximately 168 kDa and 130 kDa, respectively). Our western blot analysis revealed that Ad-ROCK1-transfected myotubes had significantly higher levels of active ROCK1 (130 kD) than those transfected with the control Ad-β-galactosidase, indicating that overexpression of intact ROCK1 resulted in the constitutive activation of ROCK1 (Figure 3a). Concurrently, we observed significant levels of mitochondrial fission in myotubes overexpressing ROCK1 compared to control C2C12 cells overexpressing β-galactosidase (Figure 3b and c). This response was associated with an increased release of ROS and cellular protein carbonylation (Figure 3d and e). Interestingly, the antioxidant N-acetyl cysteine (NAC) could abolish the protein carbonylation induced by ROCK1 overexpression, but it had no effect on mitochondrial morphology. In contrast, the addition of hydroxyfasudil (0.2 µM), a ROCK1 inhibitor, blocked mitochondrial fission (Figure 3b and c) and resulted in decreased cellular ROS levels and oxidant damage caused by ROCK1 overexpression (Figure 3d and e).

**Fig. 3.**
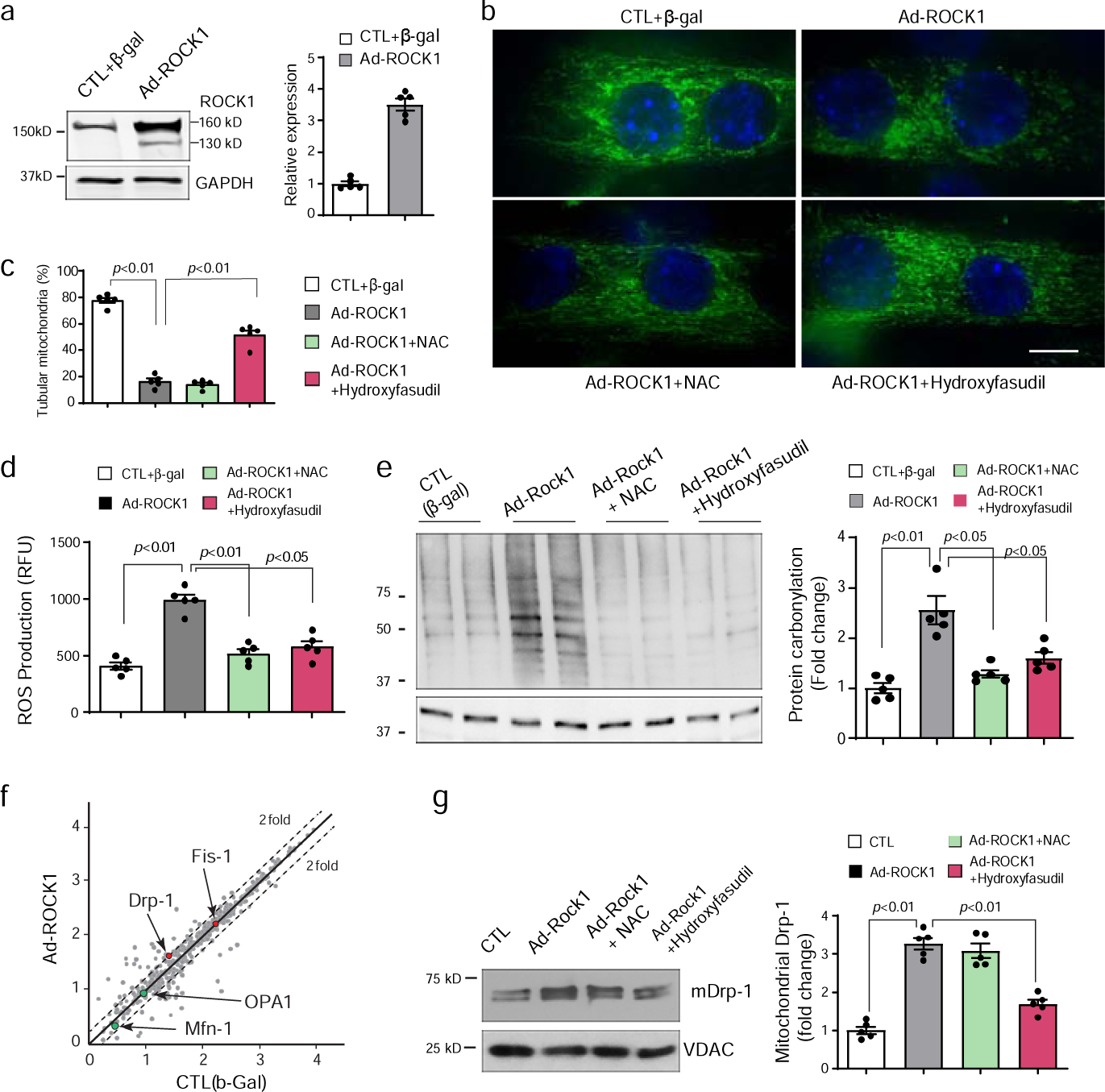
ROCK1 stimulates mitochondria fission resulting in oxidant stress in muscle cells. (A) Western blot shows increased active ROCK1(∼130 kD) protein after the overexpression of intact ROCK1 (160 kD) in C2C12 myotubes; β-gal: β-galactosidase (data are presented as mean ± SEM, n = 3). (B and C) Microscopy demonstrates fragmented mitochondria in C2C12 myotubes with ROCK1 overexpression (B). The effect of N-acetyl cysteine (NAC) and hydroxylfasudil on mitochondrial fission was also analyzed by measuring the percentage of tubular mitochondrial intensity relative to the total mitochondrial intensity (C). Scale bar = 10µm. (D) ROS production was measured in C2C12 myotubes under conditions as described (Data are presented as mean ± SEM, n = 3). (E) Representative western blot shows the levels carbonyl protein in C2C12 myotubes with treatments as described (data are presented as mean ± SEM, n=7). (F) Scattered plot depicts the mitochondrial protein content analyzed by mass spectrometry. Mitochondria fractions were isolated from myotubes with or without ROCK1 overexpression. Dash lines indicate 2-fold changes in comparison to the result of control myotubes. (G) Representative western blot shows the expression of Drp-1 in mitochondrial fraction of C2C12 myotubes. Voltage-dependent anion channel (VDAC) was used as mitochondrial protein loading control (data are presented as mean ± SEM, n=7).

Subsequently, we investigated the signaling pathway involved in ROCK1-induced mitochondrial fission. We isolated mitochondria and analyzed changes in mitochondrial protein expression in response to ROCK1 overexpression. Using protein mass spectrometry analysis, we identified 48 upregulated and 39 downregulated mitochondrial proteins during ROCK1-induced mitochondrial fractionation. Among the proteins related to mitochondrial fusion and fission, Drp1 was found to be enriched in the mitochondrial fraction from cells overexpressing ROCK1, whereas no changes were observed for proteins controlling mitochondrial fusion (Figure 3f). Further confirmation was obtained through western blot analysis, which revealed that Drp1 levels were increased three-fold in mitochondrial fractions from cells overexpressing ROCK1, and this response was inhibited by the addition of hydroxyfasudil (Figure 3g). These findings demonstrate that ROCK1 activates Drp1, which induces excessive mitochondrial fission, leading to increased ROS emission and oxidative damage in muscle cells.

### Constitutive activation of ROCK1 in muscles aggravates CKD-induced atrophy via mitochondrial fragmentation

Previously, we generated muscle-specific, constitutively activated ROCK1 (caROCK1)-expressing mice by crossing muscle creatine kinase promoter-driven Cre (MCK-Cre) mice with transgenic mice bearing a floxed, transcriptional stop cassette upstream of the constitutively active ROCK1 (caROCK1 ^f/f^). Our findings indicated that caROCK1 caused a decrease in mitochondrial content ^26^. However, using MCK-Cre to generate caROCK1 mice can lead to unpredictable developmental problems or genetic compensation in skeletal muscle. To avoid these potential issues, we developed an inducible ROCK1 activation mouse model that enabled us to activate ROCK1 specifically in the muscles of adult mice. We crossed caROCK1 ^f/f^ mice with transgenic mice carrying a doxycycline-inducible Cre recombinase driven by the human skeletal actin (HSA) promoter, resulting in HSA-Cre/ caROCK1 ^f/f^ double-positive mice ^27^. After establishing the CKD model, we administered doxycycline to the mice for two weeks to induce constitutive activation of ROCK1 in muscle, referred to as mROCK1ca mice and the activation of ROCK1 was confirmed by activity assay (Figure 4a). caROCK1 ^f/f^ mice that received doxycycline administration served as controls. Under normal conditions, the 2-week activation of ROCK1 did not cause any detectable changes in muscle architecture or myopathic features in mROCK1^ca^ mice. However, under CKD conditions, mROCK1^ca^ mice exhibited more severe muscle atrophy compared to control mice with CKD (Figure 4b). Specifically, the TA muscle mass decreased by 24% in control plus CKD mice compared to control mice with sham-operation, while mROCK1^ca^ plus CKD mice showed a 32% decrease in muscle mass compared to sham-operated mROCK1^ca^ mice (Figure 4c). The distribution of cross-sectional areas showed a clear left shift for the TA muscles of mROCK1^ca^ mice with CKD (Figure 4d). Electron microscopy confirmed the presence of excessive mitochondrial fragmentation in the muscles of mROCK1^ca^ plus CKD mice, characterized by short, swollen mitochondria and loose, disrupted myofilaments (Figure 4e). The excessive mitochondrial fragmentation observed in the muscles of mROCK1^ca^ mice was accompanied by decreased expression of OXPHOS enzymes and elevated levels of cellular protein carbonylation, indicating exacerbated oxidative stress responses and accelerated muscle protein loss in mROCK1^ca^ mice under CKD conditions (Figure 4f and g). Thus, constitutive activation of ROCK1 promotes excessive mitochondrial fragmentation in the muscle, leading to increased oxidative stress responses and accelerated muscle protein loss in mice with CKD.

**Fig. 4.**
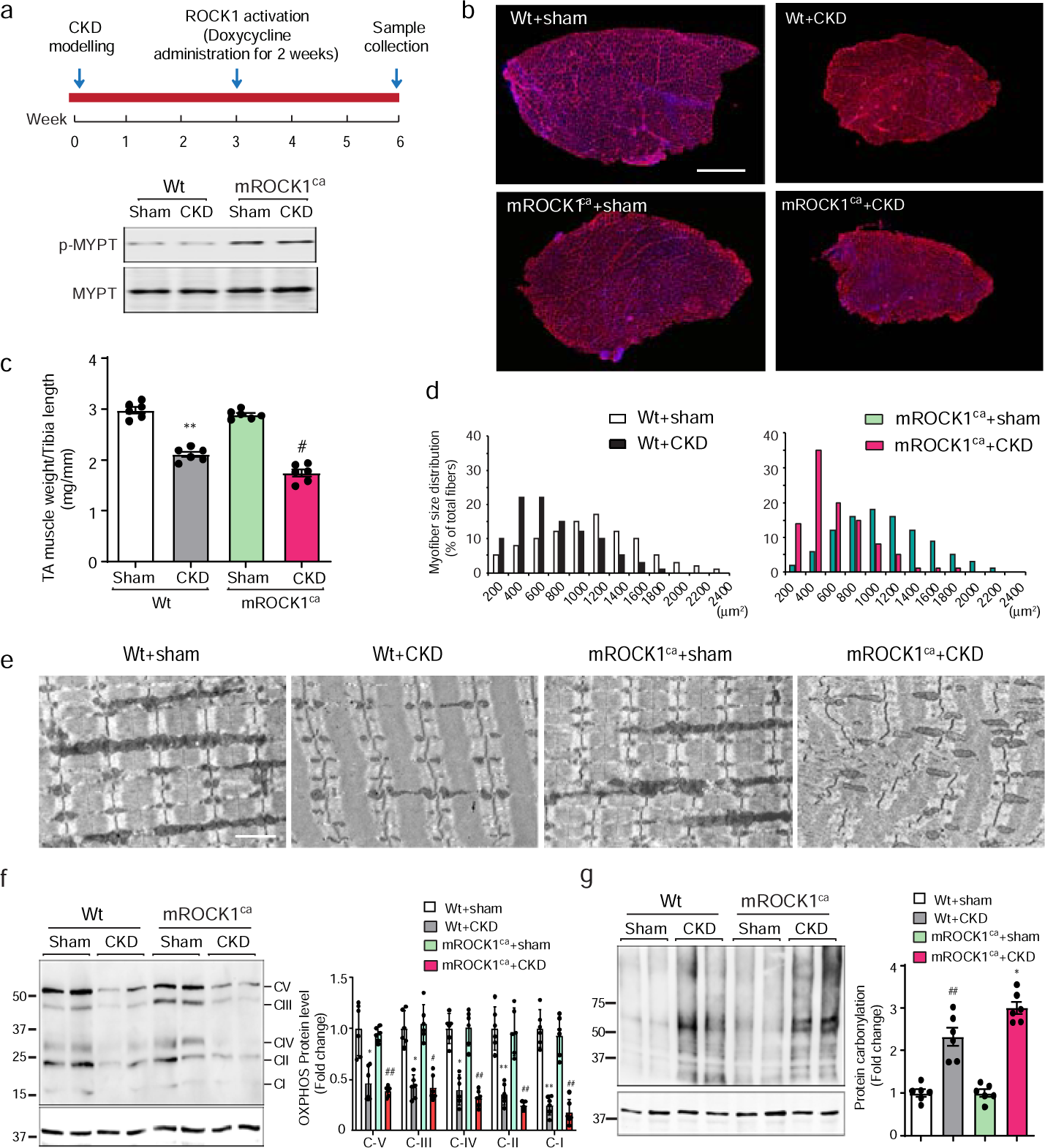
Muscle-specific ROCK1 activation aggravates muscle atrophy in vivo. (A) Upper panel: Timeline of experimental design. Lower panel: ROCK1 activity assay shows that the phosphorylation of MYPT (a substrate of ROCK1) increased in muscles of mice with muscle specific ROCK1 overexpression (mROCK1ca). (B) Immunostaining of cross sections of TA muscles from mROCK1ca mouse with or without CKD. Red represents laminin staining. Scale bar= 500 µm. (C) The quantification of TA muscle weight to tibia length ratio (n=7). (D) The distribution of myofiber cross-sectional area in TA muscles from experimental mice as indicated (n = 5). (E) Transmission electron microscopy demonstrates swelled and fragmented mitochondria in sagittal sections of muscles from experimental mice as indicated (Scale bar = 500nm). (F) Representative western blot shows the expression of subunits of mitochondrial respiratory electron transport chain in muscles from wild-type and mROCK1ca mice with or without CKD (n=7). (G) Representative western blot demonstrates carbonyl protein levels of muscles from wild-type and mROCK1ca mouse with or without CKD (n=7). All data are presented as means ± SEM (\**p* < 0.05, \*\**p* < 0.01, Wt+CKD vs. Wt+sham. #*p* < 0.05, mROCK1^ca^+CKD vs. Wt+CKD).

### ROCK1 deletion mitigates CKD-induced muscle atrophy by suppressing mitochondrial fission and oxidative damage

To verify that ROCK1 activation contributes to muscle atrophy through inducing mitochondrial fragmentation, we also investigated whether deletion of ROCK1 (ROCK1^-/-^) can attenuate muscle protein loss in mice with CKD. After confirming the absence of ROCK1 protein in muscle from ROCK1^-/-^ mice (Figure 5a), we divided the mice into four groups: wild-type plus sham, wild-type plus CKD, ROCK1^-/-^ plus sham, and ROCK1^-/-^ plus CKD. We measured muscle atrophy parameters, including the TA muscle weight/tibia length ratio and cross-sectional area distribution. We observed no significant differences in muscle atrophy parameters between sham-operated ROCK1^-/-^ mice and their wild-type littermates. However, ROCK1^-/-^ plus CKD mice displayed a marked increase in muscle mass compared to wild-type plus CKD mice (Figure 5b and c). Along with the observed improvements in atrophy parameters, electron microscopy revealed that mitochondrial damage was partially restored in ROCK1^-/-^ plus CKD mice, indicating that ROCK1 deletion suppressed mitochondrial fragmentation in the atrophic myofibers of CKD mice (Figure 5d). Mechanistically, the improvement in mitochondrial fragmentation was likely due to the inhibition of Drp-1 activity because there was a significant decrease in the quantity of Drp-1 in the mitochondrial fractions of muscles from ROCK1^-/-^ plus CKD mice compared with that in wild-type plus CKD mice (Figure 5e). Furthermore, the protein carbonylation levels in muscle from ROCK1^-/-^ plus CKD mice were significantly lower than those in wild-type plus CKD mice (Figure 5f), indicating the attenuation of oxidative stress in muscles from ROCK1^-/-^ mice. Interestingly, the observed decrease in mitochondrial respiratory electron transport protein levels in muscle from CKD mice was also partially rescued by the ROCK1 knockout, suggesting that the suppression of excessive mitochondrial fission can relieve mitochondrial damage caused by CKD (Figure 5g). Taken together, these results demonstrated that the deletion of ROCK1 prevented excessive mitochondrial fission by blocking the recruitment of Drp-1 onto mitochondria, resulting in reduced levels of oxidative damage and ameliorating muscle wasting in mice with CKD.

**Fig. 5.**
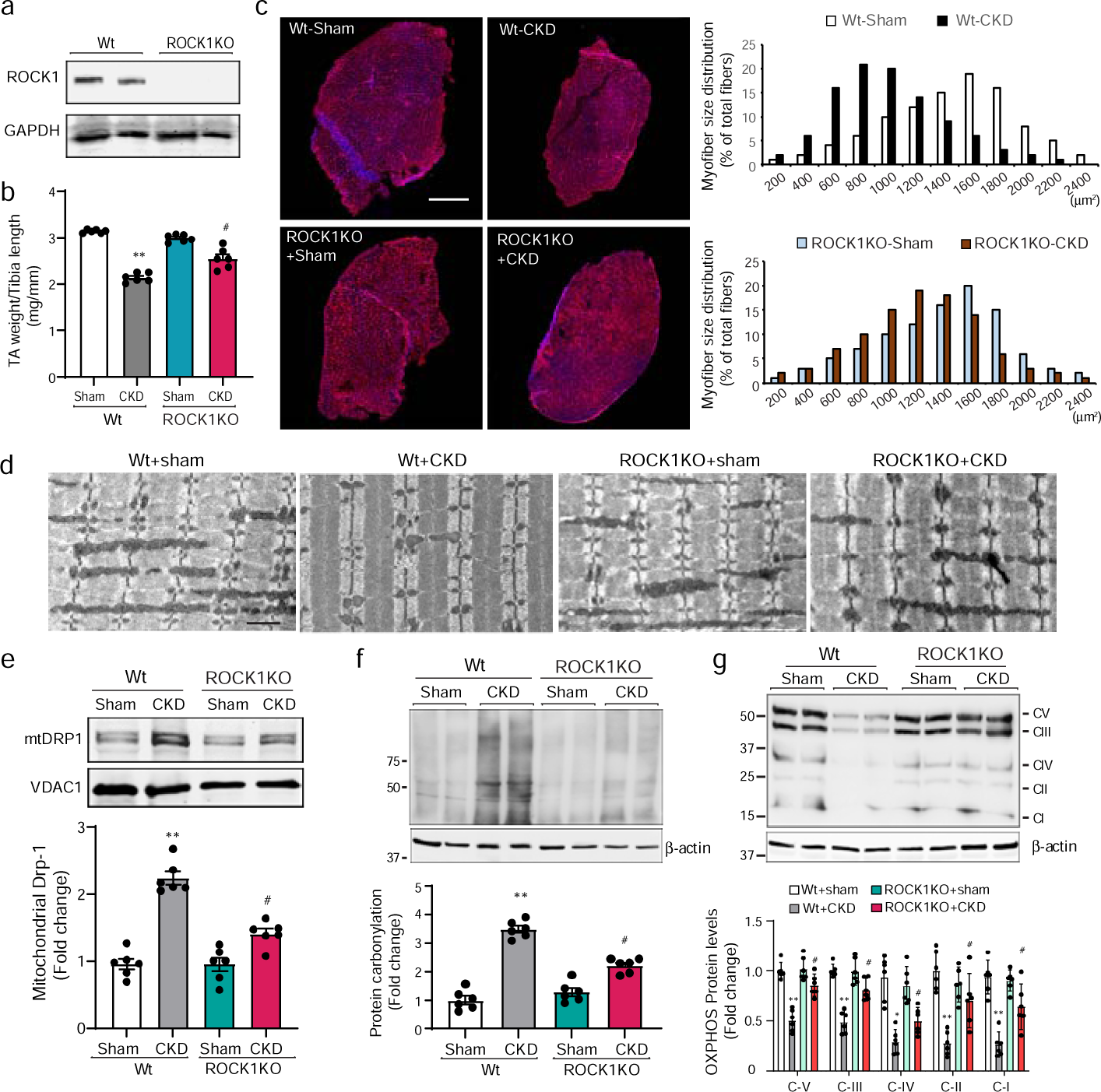
ROCK1 knockout (KO) attenuates mitochondrial fragmentation and oxidative stress in CKD mice. (A) Western blot shows the eliminated ROCK1 expression in muscles from ROCK1KO mice. (B) The quantification results of TA muscle weight to tibia length ratio (n=7). (C) Immunofluorescence using anti-dystrophin shows cross sections of tibialis anterior (TA) muscles from Wt and ROCK1 KO mice with or without CKD. Scale bar=1mm. Right panel: the leftward-shifted distribution of cross-sectional area of TA muscle induced by CKD was corrected in ROCK1KO mice despite the presence of CKD (n = 5). (D) Transmission electron microscopy demonstrates mitochondria fission caused by CKD was mitigated in Wt and ROCK1KO mice with or without CKD (Scale bar =500nm). (E) Representative western blots of mitochondrial Drp1 (mtDrp-1) from muscles of ROCK1KO mice with or without CKD (n=7). (F) Representative western blot demonstrates carbonyl protein level is suppressed in muscles from ROCK1KO-CKD mice (n=7). (G) Representative western blot shows the expression of subunits of mitochondrial respiratory electron transport chain in muscles from Wt and ROCK1KO mice with or without CKD (n=7). All data are shown as means ± SEM (\**p* < 0.05, \*\**p* < 0.01, Wt+CKD vs. Wt+sham. #*p* < 0.05, ##*p* < 0.01 ROCK1KO+CKD vs. Wt+CKD).

### Pharmacological inhibition of ROCK1 by hydroxyfasudil ameliorates muscle wasting in mice with CKD

We then investigated the potential of pharmacological inhibition of ROCK1 to attenuate muscle wasting in mice with CKD. To this end, we administered hydroxyfasudil (2 mg/kg), a selective ROCK1 inhibitor, or placebo to CKD mice and sham-operated control mice for 4 weeks. As depicted in Figure 6a, hydroxyfasudil did not affect body weight in control mice but significantly increased body weight in pair-fed CKD mice, suggesting that the drug was effective in correcting the wasting syndrome in mice with CKD. This increase in body weight was attributed to a gain in muscle mass, as evidenced by the significantly higher TA muscle weight/tibia length ratio in hydroxyfasudil-treated CKD mice compared to placebo-treated CKD mice (Figure 6b). To confirm these observations, we further analyzed cross sections of TA muscles from each group. The results showed that the average cross-sectional areas in hydroxyfasudil-treated CKD mice were significantly larger than those in placebo-treated CKD mice (Figure 6c), and an obvious rightward shift was observed in the fiber size distribution of TA muscles from hydroxyfasudil-treated CKD mice (Figure 6d). Next, we measured protein degradation rates in both the extensor digitorum longus (EDL) and soleus muscles. No differences were observed in protein degradation rates between control mice treated with placebo and control mice treated with hydroxyfasudil. However, the rate of protein degradation in muscles from CKD mice was significantly lower in mice treated with hydroxyfasudil compared to those treated with placebo (Figure 6e). This was further supported by decreased mRNA levels of MuRF-1, Atrogin-1 and MUSA1 in muscles from CKD mice treated with hydroxyfasudil compared to those treated with placebo (Figure 6f). To assess the protective effects of hydroxyfasudil against mitochondrial damage, we induced CKD in PhaM^excised^ Dentra-2 transgenic mice and treated them with hydroxyfasudil. Confocal microscopy of TA muscle revealed increased mitochondrial fragmentation in CKD mice, characterized by an increase in the number of small, punctate mitochondria in myofibers compared with control mice that had long filamentous mitochondria. However, this increased mitochondrial fragmentation was suppressed in CKD mice treated with hydroxyfasudil (Figure 6g). To quantify the changes in mitochondrial fragmentation, we plotted the results as a distribution curve ^28^. CKD mice displayed a left-shift in the mitochondrial size distribution compared with sham-operated control mice, while the size distribution of mitochondria in muscles from hydroxyfasudil-treated mice shifted significantly towards the right, indicating reduced mitochondrial fragmentation (Figure 6g, lower panel). Furthermore, the mitochondrial-associated Drp-1 level markedly decreased in mitochondrial fractions derived from muscles of hydroxyfasudil-treated CKD mice compared with that in muscles from untreated CKD mice (Figure 6h). We also evaluated whether oxidative damage could be attenuated by hydroxyfasudil administration. As shown in Figure 6i, hydroxyfasudil treatment significantly reduced the levels of protein carbonylation in muscles from CKD mice. Overall, these findings suggest that hydroxyfasudil administration attenuated excessive mitochondrial fission and ameliorated oxidative damage in the muscles of mice with CKD.

**Fig. 6.**
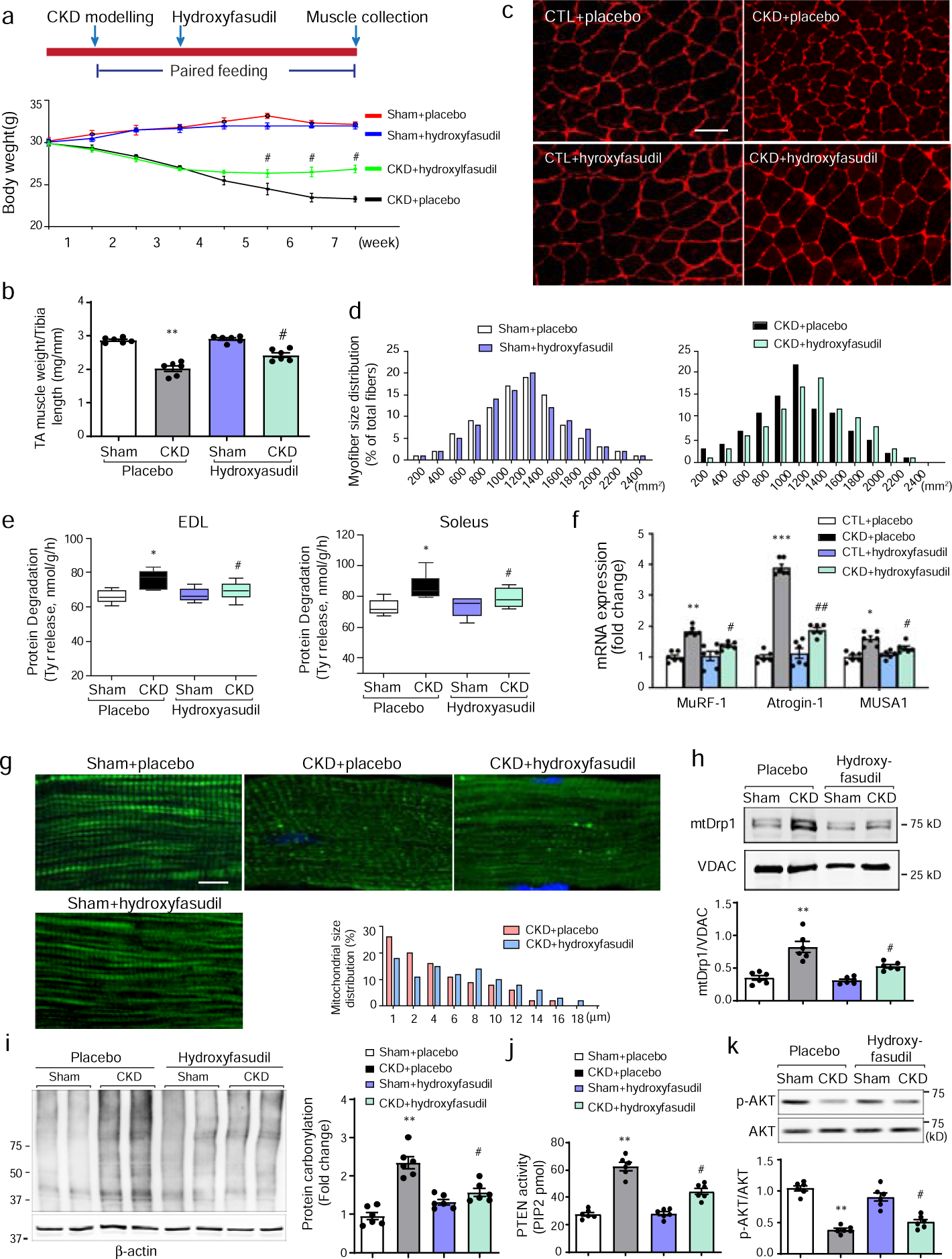
Pharmacological inhibition of ROCK1 hydroxyfasudil prevents muscle mitochondrial fission and atrophy in CKD mice. (A) Upper panel: Timeline of hydroxyfasudil administration. Lower panel: Bodyweight changes in sham-operated and CKD mice treated placebo or hydroxyfasudil (n=12). (B) The quantification of TA muscle weight to tibia length ratio (n = 7). (C) Myofibers cross sectional area in TA muscles from sham-operated and CKD mice treated with placebo or hydroxyfasudil. Scale bars: 50µm. (D) The distribution of myofiber cross-sectional area in TA muscles was shifted rightward from hydroxyfasudil-treated CKD mice compared with placebo-treated CKD mice (n = 5). (E) The rate of protein degradation of extensor digitorum longus (EDL) and soleus muscles from experimental mice (n = 7). (F) Quantitative real-time PCR shows the repressed expression of MuRF-1, Atrogin-1 and MUSA1 in the muscles from experimental mice (n=5). (G) Representative microscopy images show mitochondrial fragmentation in myofibers from CKD mice is suppressed by hydroxyfasudil administration (scale bar=10µm). Lower right panel: The distribution of mitochondrial size is shifted rightward in myofibers from hydroxyfasudil-treated CKD mice. (H) Representative western blot demonstrates Drp1 in mitochondria fractions from muscles of experimental mice (n=5). (I) Representative western blot demonstrated carbonyl protein in muscles of sham and CKD mice treated with placebo or hydroxyfasudil (n=5). (J) PTEN activity assay shows hydroxyfasudil suppressed PTEN activity in muscles of CKD mice (n=5). (K) Representative western blot demonstrates the phospho-Akt is increased by hydroxyfasudil in muscles of CKD mice (n=5). All data are shown as means ± SEM (*P < 0.05, **P < 0.01, ***P < 0.001 CKD plus placebo vs. Sham plus placebo, #P < 0.05, ##P < 0.01 CKD plus hydroxyfasudil vs. CKD plus placebo).

Our previous study indicated that ROCK1 activation leads to the suppression of Akt by activating phosphatase and tensin homolog (PTEN) ^19^. Therefore, the inhibition of ROCK1 should stimulate Akt activity. We examined the phosphorylation of Akt in muscles from hydroxyfasudil-treated CKD mice. As expected, hydroxyfasudil administration significantly suppressed PTEN activity and concomitantly increased p-Akt levels in muscles of CKD mice compared to untreated counterparts (Figure 6j and k). Thus, the observed suppression of PTEN activity in the muscles of CKD mice treated with hydroxyfasudil suggests that the drug exerts a dual effect to counteract muscle wasting by both ameliorating oxidative stress and enhancing anabolic signaling.

## Discussion

In this study, we sought to identify a novel pathway that links the dynamics of the mitochondrial network to oxidative stress and protein loss in skeletal muscle under CKD condition. Through transcriptomic analysis, we observed a simultaneous decrease in OXPHOS activity and increase in ROS emission in muscles during atrophy, representing two counteracting events. We then probed into the possibility that the activation of the mitochondrial fission machinery could be the underlying cause for the elevated ROS release and oxidative damage, which in turn might drive muscle atrophy. By employing both gain-of-function and loss-of-function approaches in vivo, we were able to elucidate the cellular signaling pathways involved in mediating mitochondrial fission during muscle atrophy induced by CKD. Specifically, we demonstrated that ROCK1 activates Dynamin-related protein 1 (Drp-1), leading to the recruitment of Drp-1 to the mitochondria. This, in turn, resulted in excessive mitochondrial fragmentation and oxidative damage in muscles. Additionally, we found that pharmacological inhibition of ROCK1 effectively suppressed mitochondrial fission in muscle of mice with CKD. These observations collectively suggest that the deleterious activation of ROCK1 and the subsequent fragmentation of mitochondria constitute a critical mechanism contributing to muscle atrophy. Our findings underscore the potential therapeutic value of targeting this pathway in the management of muscle wasting associated with CKD.

Skeletal muscle is crucial for protein and energy metabolism in mammals. Under catabolic states, a decrease in OXPHOS and changes in mitochondrial morphology are linked to muscle protein loss ^29, 30^. Conditions like cancer, CKD, diabetes, and microgravity have been associated with alterations in muscle mitochondria ^31-33^. Maintaining mitochondrial integrity is vital due to its role in limiting excessive ROS release ^34, 35^. Mitochondrial dynamics, involving continuous fission and fusion, are critical for maintaining mitochondrial morphology ^36^ and are regulated by proteins that balance these processes ^37^. Drp-1, a cytosolic GTPase, plays a central role in mitochondrial fission by partnering with FIS-1 ^38^. While fission is essential for isolating damaged mitochondria ^39^, excessive fission can lead to the loss of membrane potential, increased ROS emission, and oxidative stress ^40^. The activation of fission through the AMPK/FoxO3 pathway has been shown to cause muscle protein loss ^37^, while inhibiting fission, such as by downregulating Fis1, can prevent muscle atrophy ^41^. Given the pivotal role of mitochondrial fission in muscle catabolism, modulating the cellular signaling pathways involved in this process offers a promising approach to counter muscle loss and improve function ^42^. In this context, our study uncovers the role of ROCK1 in activating Drp-1, leading to excessive mitochondrial fragmentation and oxidative damage in muscles. By pharmacologically inhibiting ROCK1, we observed a notable suppression of mitochondrial fission in CKD mice, highlighting the detrimental effects of ROCK1 activation on mitochondrial dynamics as a key factor contributing muscle atrophy. Moreover, our investigation into mitochondrial dysfunction in CKD muscles revealed significant reductions in basal respiration, maximal respiratory capacity, reserve capacity, and maximal ATP generating capacity, as assessed through the oxygen consumption and extracellular acidification rates of freshly isolated myofibers. These reductions can be attributed to decreased OXPHOS activity and beta-oxidation capacity in atrophic muscles. Furthermore, we observed an increase in proton leak, which could potentially account for the elevated basal metabolic rates and body temperatures reported in patients with CKD, particularly considering the significant contribution of skeletal muscle to body mass ^43, 44^.

Skeletal muscle is known to produce reactive oxygen species (ROS) from various subcellular locations, but it is widely accepted that abnormal mitochondrial ROS production is the primary source triggering oxidative stress ^12^. Interestingly, as mitochondrial function decreases during muscle atrophy, ROS production should theoretically diminish. Therefore, oxidative stress likely stems not from ROS production itself, but rather from mitochondrial disruption and leakage. This insight is pivotal, as it suggests that therapeutic strategies should focus on preserving mitochondrial integrity by reducing excessive fission, rather than solely inhibiting ROS production. In this study, we addressed this by employing hydroxyfasudil, a ROCK1 inhibitor, to impede mitochondrial ROS induced cellular damage, because ROCK1 is an upstream signaling molecule implicated in promoting mitochondrial fragmentation and ROS emissions. Our results demonstrated that hydroxyfasudil effectively counteracted CKD-induced muscle atrophy by suppressing both mitochondrial fragmentation and ROS emissions. Furthermore, hydroxyfasudil elevated p-Akt levels in atrophied muscles, which is attributed to disrupting a feedback loop identified in muscles from CKD mice ^19^. This loop involves caspase-3 activating ROCK1, which in turn stimulates PTEN activity. Increased PTEN activity inhibits Akt, further amplifies caspase-3 activation, thereby increasing Atrogin-1/MuRF-1 expression and accelerating muscle proteolysis ^19^. Thus, the therapeutic inhibition of ROCK1 offers dual benefits: it alleviates mitochondrial fission and oxidative stress and prevents the suppression of Akt activity in atrophying muscles. Collectively, these effects contribute to the reduction of muscle protein loss, highlighting the clinical potential of ROCK1 inhibitors as targeted treatments for conditions characterized by muscle wasting and mitochondrial dysfunction.

In conclusion, our study uncovers a novel signaling pathway involving ROCK1-mediated activation of Drp-1 leading to mitochondrial dysfunction in CKD conditions. Based on our findings, ROCK1 inhibitors, like hydroxyfasudil, could serve as therapeutic agents to prevent muscle wasting in humans with CKD or other cachexic conditions.

## Materials and methods

### Animal models

All animal experiments and procedures were conducted in accordance with the guidelines set by the Baylor College of Medicine Institutional Animal Care and Use Committee (IACUC). Ten-week-old C57BL/6 mice or PhaM^excised^ Dentra-2 transgenic mice (Jackson Laboratory, Bar Harbor, MN) were used to create a CKD-induced muscle wasting model as previously described ^45^. Hydroxyfasudil (Cayman Chemical, Ann Arbor, MI) and water mixture were administered to CKD mice by oral gavage at a dose of 2mg/kg/day for 4 weeks. Mice with muscle-specific constitutive ROCK1 activation were generated by breeding caROCK1 ^f/f^ transgenic mice (which carry a floxed transcriptional stop cassette upstream of the constitutively active ROCK1 gene) ^21^ with mice expressing doxycycline-inducible actin promoter-driven Cre recombinase (ATCA1-Cre, Jackson Laboratory), following the breeding strategy described previously ^26, 46^. ROCK1 knockout mice were described in our previous study^19^. At the end of the experiment, the mice were euthanized, and their tibialis anterior and gastrocnemius muscles, as well as soleus and extensor digitorum longus muscle, were dissected, weighed, and either immediately examined or frozen in liquid nitrogen and stored at −80LJ.

### mRNA preparation and expression analysis

Total RNA was extracted from the whole TA muscles of control or CKD mice using Qiazol (Qiagen, Germantown, MD) and then precipitated in isopropanol. iScript cDNA Synthesis Kit (Bio-Rad, Hercules, CA) was used to synthesize cDNA for quantitative real-time PCR, which was performed using the CFX96 System (Bio-Rad) with SYBR Green. Supplementary Methods lists the primer sequences for mouse Atrogin-1, MuRF-1, MUSA, β-Actin, and RPL39. For total RNA sequencing, RNA samples that passed the quality-control examination were sent to LC Sciences LLC (Houston, TX) for library preparation and paired-end 50 bp sequencing using the standard Illumina mRNA-seq protocol (n = 3 per group). For RNA-seq analyses, the data were normalized by upper quartile normalization, and genes with sequence reads less than 3 (in at least 2 samples/each group) were filtered out from further analysis. Limma was used to perform differential gene expression analysis of RNA-seq data in each mouse model, using all genes tested for differential expression as the background. DAVID Bioinformatics Resources 6.8 (https://david.ncifcrf.gov/) was used to perform GO Biological Processes Ontology and KEGG pathway analysis. Processes and pathways with *P* < 0.05 were considered significant. GSEA was performed using a desktop application (http://www.broadinstitute.org/gsea/index.jsp) and RNA-seq data of muscles obtained from control and cachexic mice were used as two expression datasets. The gene sets were acquired from the Molecular Signatures Database (MSigDB), and the names of the gene sets used were GO OXIDATIVE PHOSPHORYLATION (M12919) and RESPONSE TO OXIDATIVE STRESS (M3223).

### Cell culture, mitochondrial staining, and western blot analyses

C2C12 myoblasts were cultured in DMEM supplemented with 10% FBS (HyClone, Logan, UT), 200 units/ml penicillin, and 50 μg/ml streptomycin (Thermo Fisher Scientific). To induce myotube formation, cells were treated with differentiation media (containing 2% horse serum) for 72 hours. Adenovirus carrying mouse ROCK1 cDNA (Vector Biolabs, Malvern, PA) was added to cells to overexpress ROCK1, and cells were cultured for 48 hours. Cells were then treated with 0.2 μM hydroxyfasudil (dissolved in DMSO) or 2 mM N-acetyl cysteine (NAC) (Sigma-Aldrich, St. Louis, MO) for 24 hours. Mitochondria were visualized by staining cells with MitoTracker Probes (Thermo Fisher Scientific) following the manufacturer’s protocol.

For western blot analysis, C2C12 cell lysates were prepared in RIPA buffer containing protease inhibitors (Roche Diagnostics, Indianapolis, IN). To examine mitochondrial-associated proteins, mitochondrial fractions were isolated using the Mitochondria Isolation Kit (Thermo Fisher Scientific) and subjected to western blotting using anti-Drp-1, VDAC antibodies, or a total OXPHOS antibody cocktail (Abcam plc, Cambridge, MA). Cellular ROS levels were measured using the Cellular ROS Assay Kit (Deep Red) (Abcam). Protein carbonylation was detected by preparing skeletal muscle lysates from approximately 50 mg of muscle by homogenizing in lysis buffer provided by the Protein Carbonyl Content Assay Kit (Abcam). The resulting supernatants were subjected to western blotting following the manufacturer’s instructions.

### Measurement of cellular respiration in fresh isolated myofibers

The intact, single fibers were isolated from rear flexor digitorum brevis (FDB) muscles of experimental mice^47, 48^. The Agilent Seahorse XFe24 Analyzer was used to determine basal oxygen consumption rates (OCR) and extracellular acidification rates (ECAR) followed by Mito-stress test (Agilent seahorse Bioscience, North Billerica, USA). Briefly, cell culture microplate (Agilent Seahorse Bioscience) was coated with Cell-Tak cell and tissue adhesive (Corning, Corning, NY) before 50 µL of 22.4 ug/mL-1 Cell-Tak solution was added into the well. After 20 min of room temperature incubation, the microplate was washed twice with sterile water. Freshly isolated FDB myofibers were resuspended in 37LJ preheated Seahorse XF Assay Medium (containing 2.5 mM D-glucose and 0.5 mM L-carnitine) and seeded onto the coated Seahorse Bioscience XFe24 cell culture microplate at a density of 100 fibers/100µL/well. A control well had 100 µL Seahorse XF Assay Medium only. After centrifuged at 200g for 1 min, a 325-µL preheated assay buffer was added to the well followed by incubation at 37°C without CO2 for 30 min. The microplate was then put into a Seahorse XFe24 analyzer and examined as previously described^49^. After measurements, assay buffer was removed from each well and added 10 µl 2x Laemmli sample buffer to completely dissolve myofibers. The yield lysate then was used to run SDS-PAGE to quantify protein content for each well.

For other experimental procedures, please refer to the Supplementary Material.

### Statistics

The data are presented as mean ± SEM and were analyzed using GraphPad Prism 7. For experiments comparing two groups, we used the Student-Newman-Kuel’s two-tailed unpaired tests. When more than two groups were compared, we used ANOVA followed by Bonferroni’s multiple comparison test to analyze differences between two interested groups. A *P*-value less than 0.05 was considered statistically significant.

## Disclosure

None of the authors have a conflict of interest with respect to this work.

## Acknowledgements

We acknowledge the National Institutes of Health Grants (5RO1-DK037175). MS was supported by Natural Science Foundation of Guangdong Province (2021A1515011780).

## Author contributions

Z.H. conceived and led this study. MS, RY, HL, FL, SJ, and HP carried out experiments, study design, and data analysis; YW, SST, FD, HP and ZH interpreted the data; and ZH wrote the manuscript. All authors had final approval of the submitted versions.

---

**Figure S1.**
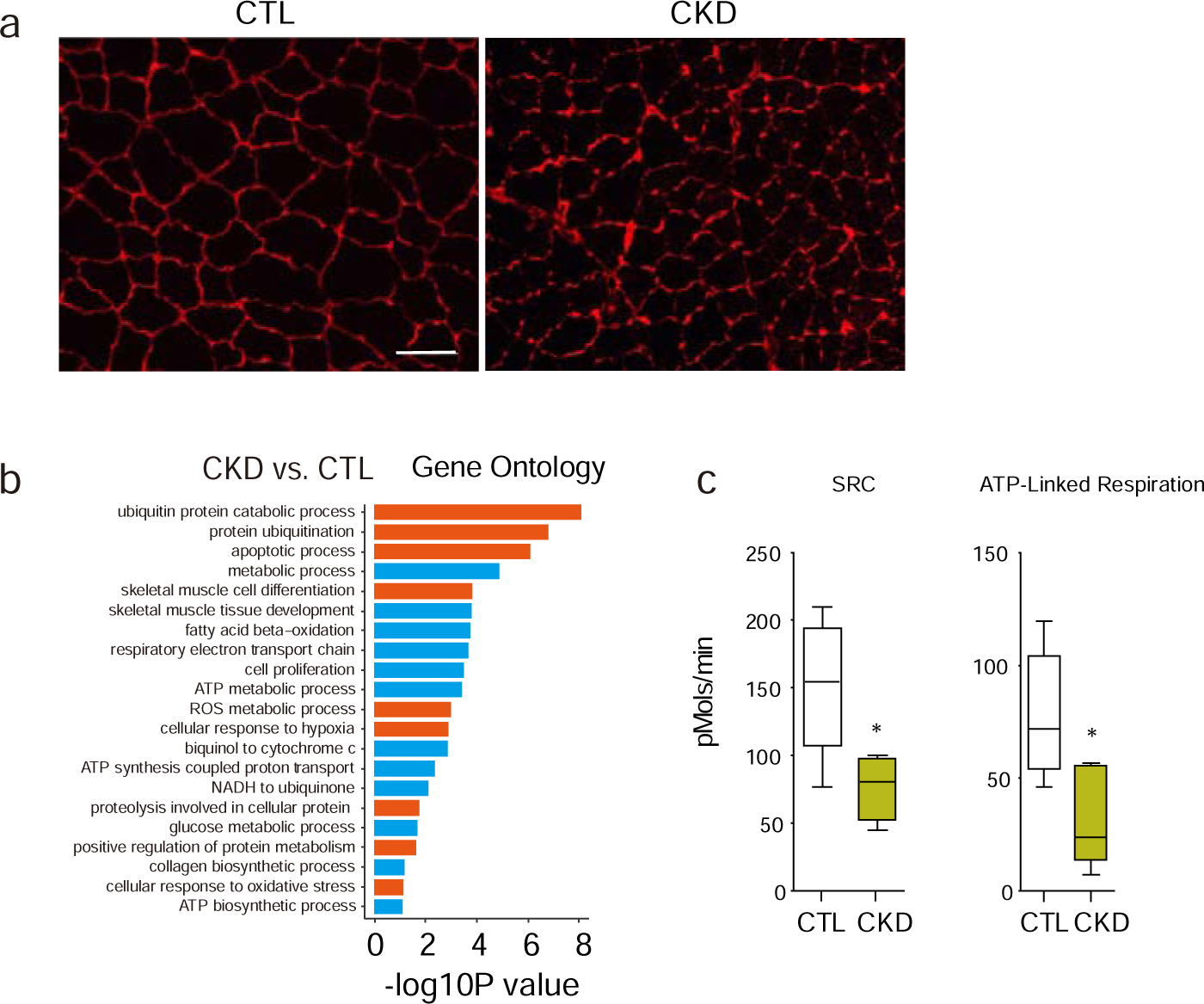
(A) Immunostaining of TA muscles from mice with chronic kidney disease (CKD, 6 weeks). Scale bars: 50 μm. (B) Gene ontology of differentially expressed genes revealed by RNA-sequence of muscles from catabolic mouse models compared with control mouse. Orange color indicates gene sets or pathways derived from genes that are significantly increased in CKD muscles, while blue indicates results analyzed from genes that are significantly decreased in CKD muscles. (C) Cellular respiration of freshly isolated rear FDB myofibers: Statistic analysis of Spare respiratory capacity (SRC) and ATP related respiration. Data are presented as mean ± SEM (\**P*<0.05, n=6).

## Notes

### Competing Interest Statement

The authors have declared no competing interest.

